# NERV: A Comprehensive Framework for Rapid, Reproducible, and Hardware-Synchronized Neuroscience Experiment Design and Execution

**DOI:** 10.1101/2025.09.04.674131

**Authors:** Kyle Coutray, Christos Constantinidis

## Abstract

**Background:** Behavioral neuroscience experiments require precise stimulus control, millisecond timing, hardware integration, and robust data provenance. Increasing use of 3D environments and multimodal recordings adds challenges for development, accessibility, and reproducibility. Fragmented tools often separate presentation, synchronization, and logging, leading to inefficiencies.

**New Method:** The Neuroscience Experimental Runtime by Vanderbilt (NERV) is a Unity-based framework that unifies experiment design, execution, and data logging. It enables rapid, no-code prototyping by automating scene and script generation, event timing, state management, hardware-synchronized data acquisition, and archival of code and experimental configurations. The modular, open-source framework implements a “low floor, high ceiling” design that lowers barriers for non-programmers while remaining extensible for advanced customization.

**Results:** Across 500 trials, Unity-to-TTL delay was 2.10 ± 1.21 ms, TTL-to-photodiode delay was 28.93 ± 0.76 ms, and Unity-to-screen delay was 31.04 ± 1.41 ms. These results confirm stable millisecond precision and frame-locked timing, enabling reliable alignment of neural, behavioral, and visual events.

**Comparison with existing methods:** Existing frameworks involve trade-offs. Some achieve precise timing but require advanced coding, while others improve accessibility but struggle with hardware or 3D graphics. Commercial platforms offer polish yet remain costly, closed-source, and inflexible. NERV combines millisecond precision, modular open-source design, and provenance in a single platform, reducing workflow fragmentation and enabling reproducible, scalable experiments.

**Conclusion:** NERV is an accessible yet extensible framework that unites rapid development, robust data provenance, and millisecond precision. It accelerates development, ensures reproducibility, and establishes a scalable foundation for next-generation neuro-science research.

## 1. Introduction

The advancement of current neuroscience research relies heavily on the development and execution of sophisticated behavioral experiments that demand precise control over stimuli, accurate timing of behavioral events relative to neural data streams, and robust data logging. Modern neuroscience experiments are increasingly complex, often requiring interactive environments, three-dimensional graphics (Brookes et al., 2020), and seamless integration with external hardware for physiological recordings such as electroencephalography (EEG) (Krugliak and Clarke, 2022), high-density microelectrode arrays e.g. Neuropixels (Jun et al., 2017), or eye-tracking systems (Iwama et al., 2024; Razavi et al., 2022). These intricate experimental designs present significant challenges in terms of development time, ease of use for researchers without extensive programming backgrounds, and ensuring the reproducibility and provenance of collected data (Davidson and Freire, 2008). Researchers frequently face obstacles in rapidly prototyping new ideas, managing diverse hardware interfaces, and archiving experimental conditions in a transparent manner for future verification and replication.

Existing tools for behavioral experiment design each offer distinct advantages but also come with notable limitations that can hinder the development of advanced neuroscience experiments. MATLAB-based Psychtoolbox (Brainard, 1997; Pelli, 1997), while celebrated as a gold standard for its precise control over stimulus timing, necessitates significant programming expertise and lacks an integrated graphical user interface (GUI) or built-in data provenance features. PsychoPy (Peirce et al., 2019) and OpenSesame (Mathôt et al., 2012), open-source Python tools, provide more user-friendly GUI builders that lower coding barriers. However, their capabilities can be limited when interfacing with complex hardware, handling graphically intensive tasks (such as 3D simulations or gamified experiments), or extending beyond simple 2D stimuli and plugins. Commercial alternatives like E-Prime (Schneider et al., 2002) or Presentation (Neurobehavioral Systems, Inc., 2025) software offer polished interfaces and reliable timing, but they are closed-source, can be costly to license, and provide limited options for customization or extension. Furthermore, many traditional frameworks require researchers to juggle multiple disparate systems, such as separate tools for stimulus presentation, hardware triggering, and data collection. This fragmentation often leads to workflow inefficiencies and synchronization errors during data merging. Few existing solutions offer the automatic archival of exact code, configuration files, and visual records that are crucial for robust data provenance (Sandve et al., 2013). To address these gaps, we developed the Neuroscience Experimental Runtime by Vanderbilt (NERV), a Unity-based framework that centralizes experimental control, hardware integration, and data logging within a single platform. NERV lowers the technical barrier to experiment design while preserving extensibility for advanced users, enabling both rapid prototyping and reproducible execution of complex behavioral tasks. We have deployed NERV in both non-human primate experiments involving high-density microelectrode recordings (similar to Riley et al. (2018)) and human studies using stereo-EEG recordings (similar to Singh et al. (2024)), where it improved experimental flexibility and integration compared to existing software used in our laboratory. In the following sections, we present the design and implementation of NERV, detailing its system architecture, user workflows, data management, hardware integration, and example experimental tasks (Section 2). We then report benchmarks of development efficiency and temporal precision (Section 3). Finally, we compare NERV with existing tools, discuss current limitations, and outline future directions (Section 4).

## 2. Methods: System Design and Implementation

This section presents NERV’s design and implementation, including its overall architecture, editor and runtime components, user workflows, data management strategy, multi-display support, hardware integration, and example experimental tasks.

### 2.1. System Overview

The Neuroscience Experimental Runtime by Vanderbilt (NERV) is a self-contained Unity framework written in C#. It automates experiment assembly through GUI-driven editor tools that generate:

1. **Task-specific Unity scenes** containing a predefined dependency prefab (camera, stimulus spawner, UI, pause controller).
2. **An auto-written** `TrialManager{Acronym}.cs` **script** that implements the user’s state machine as a single, readable `C#` coroutine.
3. **Configuration comma-separated values (CSV) files** for trials and stimulus indices, parsed at runtime by a singleton `GenericConfigManager`.

These features yield two practical benefits:

- **Rapid prototyping for experimenters without extensive programming experience:** complete runnable tasks can be generated and piloted within minutes using point-and-click form-based tools.
- **Deep extensibility for programmers:** the generated `TrialManager` is intentionally transparent so that developers can insert custom logic anywhere in the state handlers without touching the underlying engine.

During execution, NERV’s runtime layer (Fig. 1) coordinates stimulus presentation and state transitions while executing and logging all events with millisecond precision. A persistent `SessionLogManager` streams these events to disk, handles TTL output via `System.IO.Ports` (to an Arduino trigger board), and interfaces with eye-tracking systems. In our current implementation, eye-tracking input is acquired through a custom National Instruments Data Acquisition (DAQ) wrapper, though the modular design allows integration with alternative hardware. Every session’s code, configuration CSVs, logs, and screenshots are automatically archived in a timestamped folder, ensuring full data provenance.

**Figure 1:**
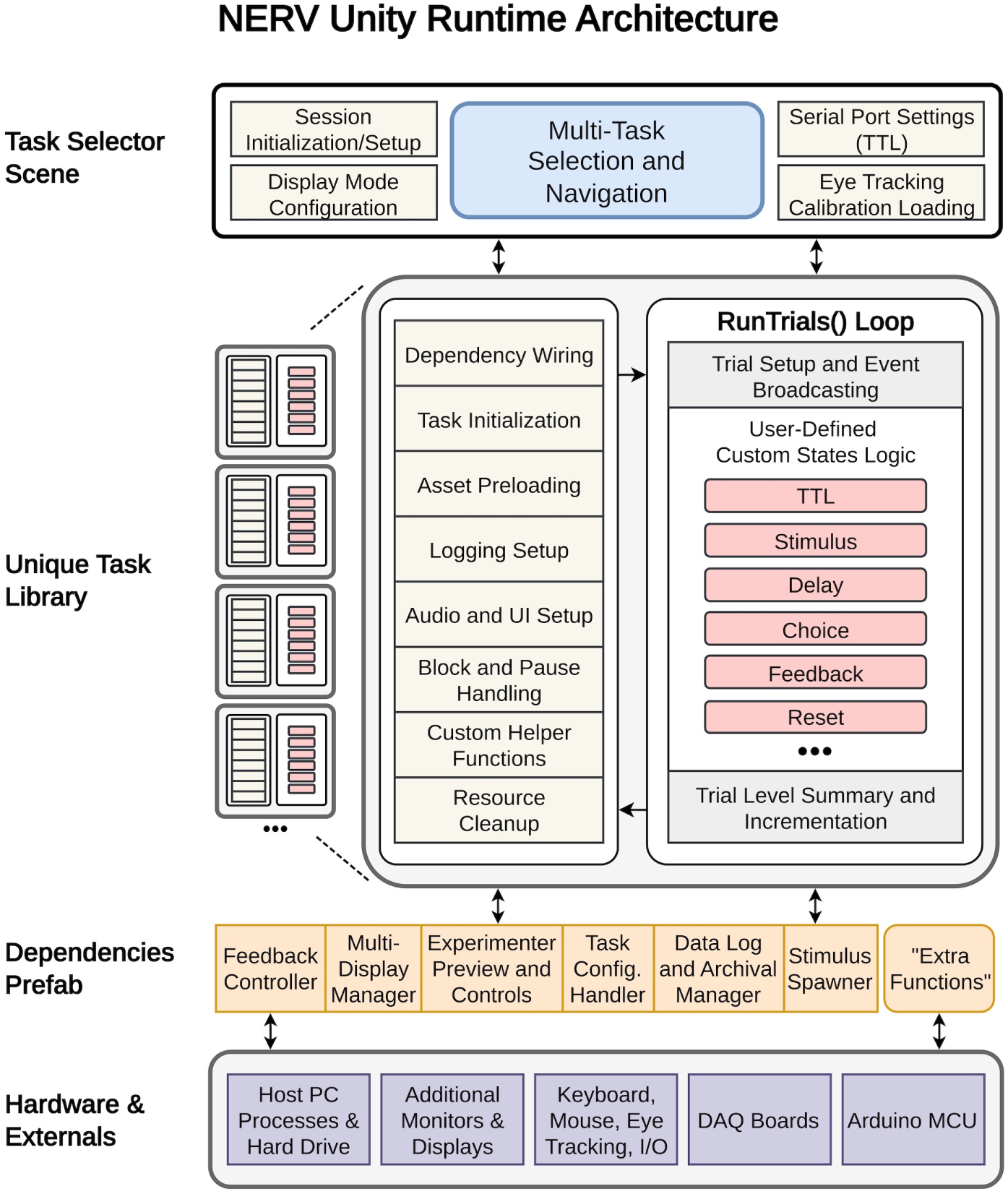
High-level overview of the NERV system architecture, showing the runtime execution layer and its connections to core components and hardware.

Together, these features establish NERV as a unified framework that couples rapid experiment generation with precise runtime control, providing the foundation for the detailed components described in the following subsections.

### 2.2. System Architecture & Core Components

The architectural backbone of NERV consists of two layers: a set of GUI-driven Editor Tools that auto-generate task assets, and a Runtime Framework of singleton `C#` components that execute, log, and synchronize the experiment. The following subsections enumerate these modules in the order they are encountered during the experiment development life-cycle.

#### 2.2.1. Editor Tools (Assets/Scripts/Editor)

##### ExperimentDefinition

A Unity `ScriptableObject` that acts as the blueprint for an experiment. Users list state names and mark their types (e.g., `IsTTL, IsStimulus, IsDelay, IsChoice, IsFeedback, IsClear`), adding parameters such as TTL codes or delay durations where relevant. The `TaskGenerator` uses this asset to create the experiment’s scene and its task-specific `TrialManager` script, while the `TrialDefinitionGenerator` references it to determine state types when generating the trial-behavior configuration file.

##### StimIndexMappingWindow.cs

An Editor Window tool responsible for generating an `{Acronym}_Stim_Index.csv` file. This CSV maps sequential numerical indices to the filenames of stimulus prefabs (e.g., .fbx files) located in a specified Unity project folder. This CSV is later parsed by `GenericConfigManager` at runtime to enable dynamic stimulus loading.

##### TrialDefinitionGeneratorWindow.cs

An Editor window that creates a *balanced* `{Acronym}_Trial_Def.csv` file. The CSV lists `TrialID` and `BlockCount`, then, for each `IsStimulus` state, dynamically includes arrays `[StateName]StimIndices` and `[StateName]StimLocations`. The generator:

– auto-computes `TrialsPerBlock` and `TotalTrials` from the user parameters S (stimuli set size), N (states), M (repetitions), and B (blocks), enforcing the balance constraint (*S × M*) mod *N* = 0;
– supports random stimulus and location sampling, per-state % occurrence, and stimulus repetitions for behavioral tasks;
– prepends a user-defined number of practice trials (recorded as block = 0), randomly sampled without replacement from the balanced trial pool.

##### TaskGeneratorWindow.cs

This core Editor Window tool automates the creation of a new Unity scene and its primary control script, `TrialManager{Acronym}.cs`. It dynamically generates `C#` code for `TrialManager{Acronym}.cs` based on the ExperimentDefinition asset, incorporating boilerplate code, dependencies, and logic tailored to the defined states (e.g., stimulus presentation, delays, choices, feedback, logging). It also instantiates a Dependencies prefab and configures GenericConfigManager. This tool is central to NERV’s “no-code” experiment generation.

##### BuildMirrorToStreamingAssets.cs

An Editor script (`IPreprocessBuildWithReport`) that executes automatically before each Unity build. It mirrors essential runtime files, specifically the generated `TrialManager{Acronym}.cs` scripts and experiment configuration .csv files, from their development directories into the `StreamingAssets` folder. This step ensures that all required code and configuration data are included in compiled standalone builds and remain accessible for data logging.

#### 2.2.2. Runtime Framework {Assets/Scripts/Core}

##### TrialManager{Acronym}.cs

The auto-generated, task-specific `C#` script that serves as the experiment’s master controller. It encapsulates the full finite-state machine in a single, readable coroutine, pulling trial definitions from `GenericConfigManager`, coordinating stimulus presentation and timing, driving choice/feedback logic, and invoking all core services (`SessionLogManager, BlockPauseController, CoinController`, and hardware-sync routines). Its transparent structure makes it the primary customization point for advanced users (see Section 2.3.2).

##### TrialData.cs

A serializable `C#` class that defines the data structure for a single row parsed from the trial definition CSV. It includes TrialID, BlockCount, and flexible dictionaries (`[StateName]StimIndices` and `[StateName]StimLocations`) to store state-specific parameters for dynamic experiment configurations

##### DependenciesManager.cs

A singleton class that provides a centralized point of access for critical runtime dependencies within the scene, simplifying the process of connecting core components (e.g., `PlayerCamera, StimulusSpawner, CalibrationLoader.cs,UI_Canvas, PauseController`) to the TrialManager{Acronym}.cs script

##### GenericConfigManager.cs

A singleton class responsible for parsing the experiment configuration CSVs (`{Acronym}_Trial_Def.csv` and `{Acronym}_Stim_Index.csv`) at runtime. It loads these files from the `Resources/Configs/{Acronym}` folder and makes the parsed trial data and stimulus index mappings available to other scripts.

##### SessionLogManager.cs

A persistent singleton (`DontDestroyOnLoad`) responsible for all session-level data logging and serial communication. It creates a timestamped folder hierarchy for each session, archives the exact `TrialManager` source code and configuration CSVs for full provenance, and maintains two primary log files: `ALL_LOGS.csv` (chronological behavioral events) and `TTL_LOGS.csv` (hardware-pulse events). Both files timestamp every entry with Unity’s monotonic clock (`Time.realtimeSinceStartup`) and a high-resolution `CPU Stopwatch`, yielding millisecond temporal precision. Furthermore, it spawns a non-blocking coroutine that asynchronously captures screen images to a `StatesCaptured` subfolder a specified number of frames after each state onset, up to a user-defined quota per state, thereby providing visual records without disrupting stimulus timing.

##### BlockPauseController.cs

Manages the in-game pause UI by displaying messages such as `PRACTICE, Block X of Y`, or `GAME COMPLETE`. It also controls the visibility and functionality of the screen labeled `CONTINUE`, the settings cog, audio controls, and the `Exit Session` button, allowing users to pause, resume, adjust settings, or return to the `TaskSelector` scene while ensuring the time scale is restored.

##### CoinController.cs

Implements the in-game coin feedback system, managing the visual representation of accumulated “coins” in a UI bar, animating coins, and providing visual and auditory feedback when the bar is filled or coins are removed.

##### TaskSelectorUI.cs

The main UI controller for the experiment selection screen, enabling users to input a session name, select tasks, configure settings, and start a new session. It also holds the `TaskMenuController` which populates the grid of selectable tasks dynamically, instantiating buttons for each available `ExperimentDefinition` asset.

##### DisplayManager.cs

A singleton class that manages multi-display configurations for experimental setups, allowing for dedicated subject and experimenter screens. It handles the activation of displays and the dynamic visibility of UI elements (e.g. hiding feedback/scoring text for non-human subjects) based on the selected mode (SingleHuman, DualHuman, DualMonkey).

#### 2.2.3. CSV Configuration Files Schema

NERV’s task logic is driven by two comma-separated value (CSV) tables that are auto-generated by the Editor tools yet remain human-readable for post-hoc editing or custom design. Both files are parsed at runtime by `GenericConfigManager`; together they fully specify trial structure and stimulus lookup.

##### **Trial Definition CSV (**{Acronym}_Trial_Def.csv)

Parsed by the `GenericConfigManager` and generated by the `TrialDefinitionGenerator`, this CSV defines every trial’s parameters in a tabular format. It typically includes:

**– TrialID:** A string identifying a unique trial, often prefixed (e.g., “WMTEST_001”).
**– BlockCount:** An integer representing the block number to which the trial belongs.
**– [StateName]StimIndices:** An array of integers mapping to stimulus prefab filenames.
**– [StateName]StimLocations:** An array of `Vector3` coordinates for stimulus placement (See Section 2.2.4).

*Note: Users may also manually edit this CSV to fine-tune trial parameters or add custom logic*.

##### Stimulus Index Mapping CSV ({Acronym}_Stim_Index.csv)

Parsed by the `GenericConfigManager` and generated by the `Stim Index Mapping` tool, this CSV maps sequential numerical indices to the filenames of stimulus prefabs (e.g., `.fbx` files) located in a specified Unity project folder. It typically includes:

**– Path to the Stimuli folder:** A string in the first line of the CSV representing the path to the folder containing stimulus prefabs. The default path is `Assets/Resources/Stimuli/Default`.
**– Index:** An integer representing the sequential index of the stimulus.
**– Filename:** A string representing the filename of the stimulus prefab.

#### 2.2.4. World Space Locations for 2D Task Development

In NERV, 2D tasks are implemented within a 3D world space, enabling flexible stimulus placement and precise control over visual presentation. Stimuli are positioned using world-space coordinates specified in the Trial Definition CSV file. The `GenericConfigManager` interprets these coordinates and communicates with the `StimulusSpawner` to place each stimulus in the Unity scene.

At a depth of *Z* = 0, world coordinates span approximately *±*4 along the horizontal (X) axis and *±*2 along the vertical (Y) axis, corresponding to the edges of the camera’s orthographic field of view. This configuration enables precise specification of spatial layouts and supports experiments requiring controlled arrangements or interactions between stimuli. By leveraging Unity’s 3D rendering environment, NERV allows 2D tasks to incorporate depth cues and realistic visual features while maintaining strict experimental control. A diagram of all world positions is shown below (Figure 2).

**Figure 2:**
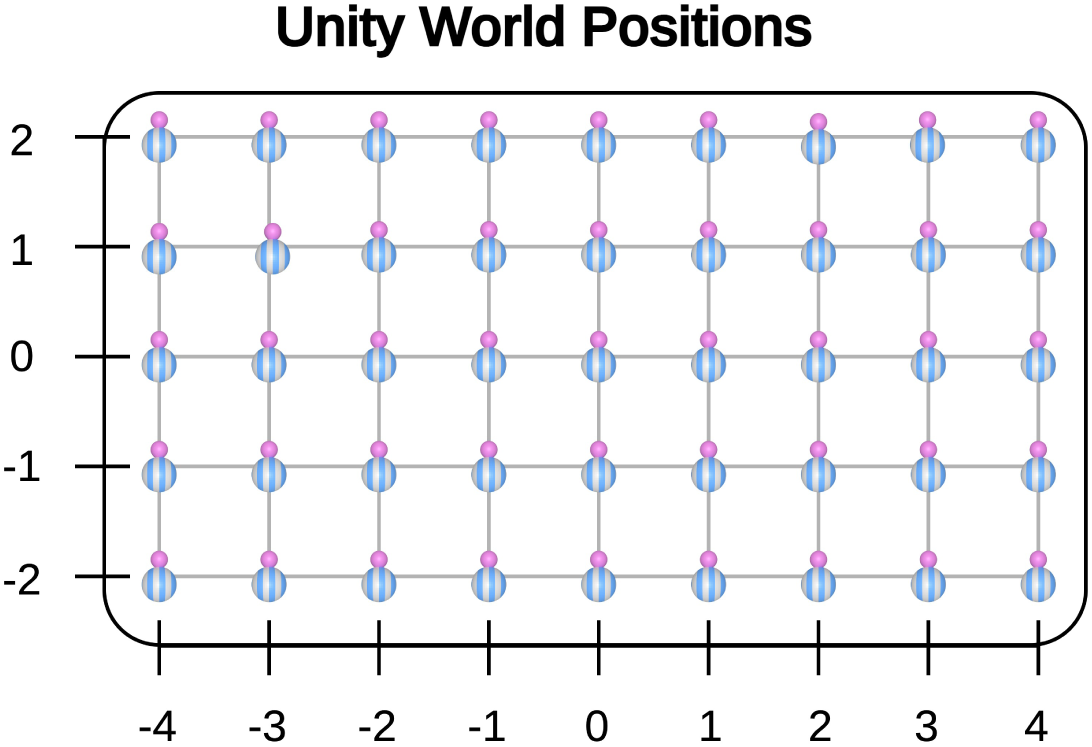
NERV’s world space coordinate system for 2D task development. Stimuli are positioned using world space coordinates, allowing for precise control over their placement within the Unity scene. The diagram illustrates the coordinate ranges and their correspondence to the camera’s orthographic view.

#### 2.2.5. NERV Default Stimulus Prefabs

NERV includes two sets of default stimulus prefabs, stored in `Assets/Resources/Stimuli`, to support rapid prototyping and testing. The current sets are based on the Quaddle stimulus library described by Watson et al. (2019), comprising 24 unique objects defined by combinations of shapes, colors, and textures. These prefabs can be instantiated and manipulated directly within the Unity environment. Although Quaddles are provided as convenient examples, users may generate and import custom prefabs to accommodate specific experimental requirements.

**Figure 3:**
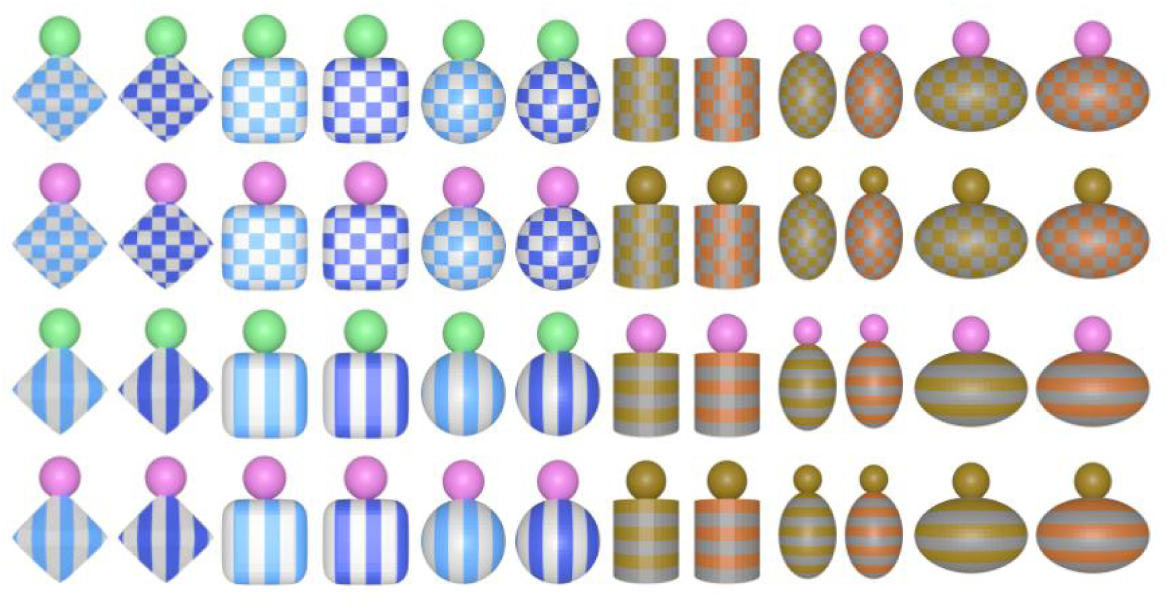
NERV’s default stimulus prefabs located in Assets/Resources/Stimuli. These prefabs provide a variety of shapes, colors, and textures for rapid prototyping and testing of experiments.

#### 2.2.6. Drag-and-Drop ExtraFunctions Modularity

Beyond its core state machine logic, the auto-generated `TrialManager{Acronym}.cs` script is designed with a powerful, decoupled extensibility point. It systematically broadcasts all experimental log events (e.g., state transitions, user inputs) via Unity’s BroadcastMessage function. This design enables the integration of what we term **ExtraFunctions**-independent Unity components (such as `FixationDotSpawner.cs` and `PhotodiodeMarker.cs`) that can be simply added via drag-and-drop to the TrialManager GameObject or its hierarchy in the Unity Editor. These ExtraFunctions (shown in Figure 4) listen for OnLogEvent calls and react to task events without requiring any direct modification of the TrialManager’s `C#` source code. This provides a robust mechanism for adding specialized experimental tools, custom logging behaviors, or hardware integrations in a modular and non-intrusive way. ExtraFunctions (located in `Assets/Scripts/ExtraFunctions`) are designed for extensibility, allowing researchers to implement additional features such as visual markers, custom feedback mechanisms, or specialized data logging formats.

**Figure 4:**
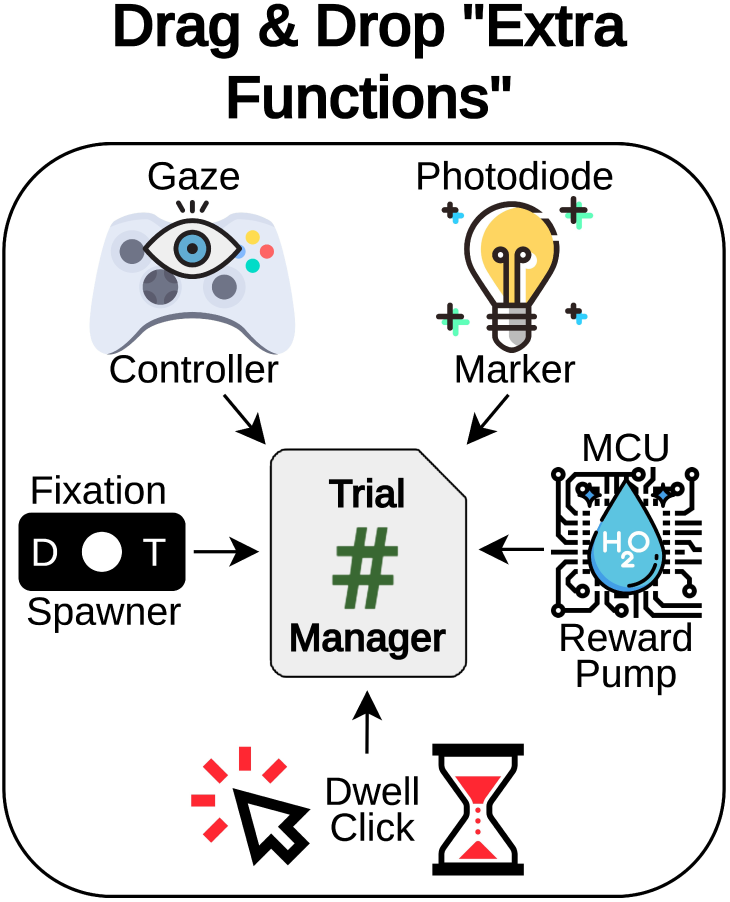
NERV’s ExtraFunctions modularity allows users to enhance their experiments by simply adding pre-built components to the TrialManager GameObject. These components listen for log events and can implement custom behaviors without modifying the core TrialManager code.

**ExtraFunctions Available in the Current NERV Version:**

1. **FixationDotSpawner.cs:** Spawns a white fixation dot in the middle of the screen during user-defined states. Users can configure the dot’s size, color, and duration in the inspector if needed, though default values are provided. To add states where the dot should appear, simply press the “+” mark in the inspector of the attached script, and type the state name exactly as defined in your TrialManager script.
2. **PhotodiodeMarker.cs:** Presents a white square in the upper-left corner at user-defined state onsets, enabling empirical measurement of code-to-screen latency with an external photodiode setup (see Section 3.1). The square’s visibility is toggled in the same frame that the stimuli appear, allowing frame-accurate latency measurements, and the flash duration can be configured in the inspector. Similarly to `FixationDotSpawner.cs`, users can add states where the marker should appear by pressing the “+” mark in the inspector and typing the state name exactly as defined in the TrialManager script.
3. **DwellClick.cs:** Enables “dwell-to-click” behavior on any stimulus object (i.e., any collider bearing a `StimulusID` component). In the inspector, users can configure `dwellTime` (the hover duration required to register a click) and a `simulateWithMouse` toggle that allows mouse-based testing of gaze behavior. When the same `StimulusID` is targeted continuously for the specified `dwellTime`, the script raises `DwellClick.ClickDownThisFrame`, which registers as a “Click” in the `TrialManager` during `IsChoice` states. Users may also choose to visualize the gaze point with a `GazeCursor` toggle in the inspector.; For genuine eye-tracking integration, the script determines a variable screenPos each frame and casts a ray through that point, using either `Input.mousePosition`, the output of an eye tracker, or the world-space location of the `GazeCursor` (controlled by `GazeCursorController`). The source code includes placeholders for SDK integration (e.g., Tobii, EyeLink, Pupil Labs), enabling straightforward extension to specific hardware APIs.
4. **GazeCursorController.cs:** Controls the position of the `GazeCursor` using data from an eye tracker. In the current implementation, this is provided through a National Instruments DAQ device, but the design can be adapted to any eye-tracking API. It works in conjunction with `CalibrationLoader.cs` to load an eye calibration JSON file and apply it to the gaze cursor’s position. Together with `DwellClick.cs`, these scripts enable gaze-based interaction in experiments. The `GazeCursorController` can also simulate NI DAQ input to verify experimental flow and calibration loading without requiring a physical eye tracker. For steps to integrate additional hardware with NERV, see Section 2.6.2.
5. **RewardPump.cs:** Enables integration of an Arduino-based reward pump, allowing experimental designs to incorporate precise, programmable reward delivery for non-human subjects. Experimenters can configure the number of bursts, the triggering state, and the duration of each burst directly in the inspector. In the current implementation, the pump is controlled via a National Instruments DAQ device interfaced with an Arduino, but the script can be adapted to operate over standard serial communication. Details of our reference hardware implementation, including build instructions and additional resources, are provided in the supplemental material.

### 2.3. User Workflow for Experiment Development

NERV provides two complementary approaches for experiment development: a no-code workflow for rapid task assembly and a scripting workflow for advanced customization.

#### 2.3.1. No-Code Pathway

This workflow, designed for users with minimal programming experience, enables the creation of complex experiments through a form-based graphical interface.

##### Step 1: Define Experiment with ExperimentDefinition

- Users create an `ExperimentDefinition` asset by right-clicking in the Project window, selecting Create *→* Experiment Definition, and assigning a custom name.
- In the Inspector for that asset, users define:

– A unique acronym (e.g., three characters).
– Each state (e.g., “Sample”, “Delay”, “Choice”, “Feedback”).
– State types via checkboxes (`IsTTL, IsStimulus, IsDelay, IsChoice, IsFeedback, IsClear`).
– For `IsTTL` states, assign a TTL code number (1–7; code 8 is reserved for `StartEndBlock`) (See Section 2.6.1 for details on how TTL states are encoded and sent to the Arduino-based interface; build instructions in Appelhoff and Stenner (2021)).
– For `IsDelay` states, input the delay duration (seconds).
- This asset serves as the blueprint for the experiment structure.

##### Step 2: Generate Stimulus Mappings with StimIndexMappingWindow.cs

- Users open Tools *→* Stim Index Mapping in the Unity menu.
- In the window:

– Enter the experiment acronym.
– Specify the Stimuli Prefab Folder (e.g., `Assets/Resources/Stimuli/Default`).
– Confirm the Output Config Folder (`Assets/Resources/Configs/{Acronym}`).
– Click Generate CSV.
- This produces `{Acronym}_Stim_Index.csv`, mapping indices to stimulus prefabs.

##### Step 3: Define Trial Sequences with TrialDefinitionGeneratorWindow.cs

- Users open Tools *→* Trial Definition Generator.
- In the window:

– Drag the `ExperimentDefinition` asset into the Experiment Definition field.
– Drag `{Acronym}_Stim_Index.csv` into the `StimIndex CSV` field.
– Input basic parameters:

* `TrialID Prefix` (e.g., `WMTEST`).
* Total Trials.
* Number of Blocks.
– For each stimulus state, configure:

* `Is Cue` toggle and (if enabled) set `Cue Repetitions` and `# Cue Stimuli per Trial`.
* `% Occurrence` (100% to always, 50% for half).
* `Random Stimuli` toggle: if enabled, specify `# Stimuli`; otherwise, list `Custom Stim Indices`.
* `Random Locations` toggle: if enabled, define Min/Max (X,Y); otherwise, list `Custom Locs` (e.g., `x,y,z;x2,y2,z2`).
– Click Generate CSV to produce {Acronym}_Trial_Def.csv.

*Note: when a state is marked `IsCue`, the generator links its indices to the final stimulus state automatically (see Automatic Cue-Target Synchronization below)*.

*Automatic Cue-Target Synchronization.* Marking a state `IsCue` triggers an internal routine that propagates the cue’s stimulus indices into the final `IsStimulus` state, enabling the default choice handler to recognize any of those indices as a correct target:

– During the `IsCue` state, the system caches the cue indices in cueIdxs[].
– When the generator builds the final `IsStimulus` state it *prepends cue indices up to the state’s capacity*, then fills any remaining slots with the state’s usual random-or-custom selection.

* **Last-state capacity** *≥* **#cue indices**: all cue items re-appear, followed by additional (random) stimuli.
* **Last-state capacity** *<* **#cue indices**: only the first *N* cue indices (*N* = capacity) are inserted.
– At choice time the `TrialManager` scores `correct = answered && cueIdxs.Contains(pickedIdx);`

i.e. any stimulus that was presented in the cue epoch is considered a correct selection.

##### Step 4: Generate Task with TaskGeneratorWindow.cs

- Users open Tools *→* Task Generator.
- In the `Task Generator` window:

– Drag the `ExperimentDefinition` asset into the `Experiment Definition` field.
– Drag the NERV `Dependencies` prefab into the `Dependencies Prefab` field.
– Click `Generate TrialManager`.
- After generation, a new button Attach TrialManager Now appears.
- Users click `Attach TrialManager` Now to attach the `TrialManager{Acronym}.cs` script to the scene and wire all core dependencies.
- Once attached, the tool will have automatically:

– Created a new Unity scene named after the experiment.
– Instantiated the `Dependencies` prefab and connected components (camera, stimulus spawner, UI, pause controller).
– Configured `GenericConfigManager` for runtime CSV loading.
- The experiment is now fully assembled and ready to run. Users press `Play` or `Build and Run` to pilot the task—complete with stimuli, timing, input handling, and data logging—without writing code. With proper planning, this rapid prototyping workflow enables a complete design-to-execution cycle in just minutes to hours. A simplified overview of the no-code workflow is shown in Figure 5.

In summary, the no-code pathway allows researchers to assemble and pilot complete experiments with minimal technical overhead, making it well suited for rapid prototyping and behavioral testing.

**Figure 5:**
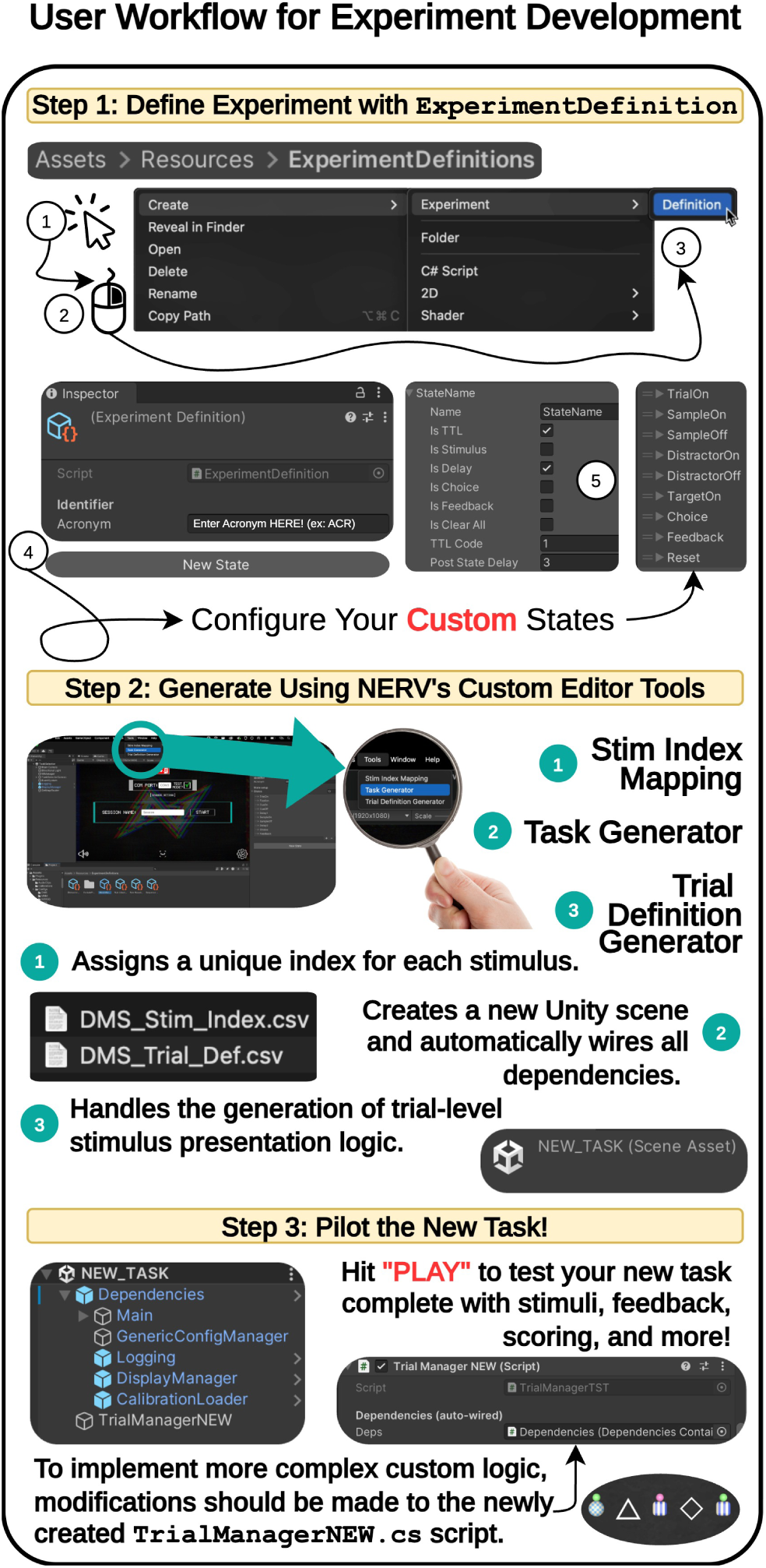
Simplified overview of NERV’s no-code experiment development workflow. Users define the experiment structure, configure stimulus mappings and trial sequences, and build the complete task scene and controller script—all through intuitive GUI tools without writing code.

#### 2.3.2. **Flexible Coding Pathway:** Customizing the TrialManager Script

For users with programming skills or specialized requirements, NERV extends beyond the no-code workflow to support deeper customization. At the center of this pathway is the `TrialManager{Acronym}.cs` script, which governs the experiment loop and manages all trial-specific behaviors and state transitions. Although it is automatically generated by the `TaskGeneratorWindow` from the user’s `ExperimentDefinition`, the script is intentionally kept human-readable and well-commented so that developers can easily adapt it to their needs.

The `TrialManager` is organized as a linear “runner” loop implemented as a `C# IEnumerator` coroutine named `RunTrials()`. This structure traces the logical flow of a trial from start to finish, with clearly demarcated sections for each user-defined state (e.g., “Stimulus,” “Delay,” “Choice,” “Feedback,” “ClearAll”). By following this layout, developers can quickly locate and modify specific behaviors to implement custom logic such as complex trial structures, dynamic stimulus generation, or advanced input handling.

Key aspects for customization include:

- **Direct Modification of State Logic:** Developers can open the `TrialManager` script and insert custom code within the pre-defined sections for each state. For example:

– Implement novel scoring algorithms or adjust existing ones based on complex criteria.
– Implement complex decision-making algorithms in the “Choice” state, allowing for adaptive trial structures.
– Create dynamic stimulus behaviors that react to real-time participant input or internal game state beyond simple presentation.
– Add specialized interactions with game objects, physics, or other Unity features that are not covered by NERV’s standard components.
– Modify feedback mechanisms to include additional visual or auditory cues based on user-defined criteria.
- **Configurable Public Variables:** The generated `TrialManager` script exposes many key parameters as public variables in the Unity Inspector. This allows researchers to adjust critical settings like `MaxChoiceResponseTime, FeedbackDuration, PointsPerCorrect, CoinsPerCorrect`, and `[StateName]Duration` for each user-specified `IsDelay` state directly within the Unity editor without needing to alter the `C#` code itself.
- **Interfacing with Core Modules:** The `TrialManager` automatically wires its necessary dependencies through the `DependenciesManager.cs`. It retrieves trial data from `GenericConfigManager.cs`, logs all events and sends TTL pulses via `SessionLogManager.cs`, manages pause screens with `BlockPauseController.cs`, and provides real-time feedback using `CoinController.cs`. This central integration simplifies customization, as coders primarily interact with the `TrialManager` script and its exposed parameters and helper functions.
- **Transparent Logic:** Unlike “black-box” graphical user interface (GUI) experiment builders that hide underlying logic, NERV’s approach provides a transparent view of the task logic. This transparency is invaluable for debugging, verifying experimental conditions, and extending functionality, giving users complete control over their experiment’s behavior.

In essence, by maintaining a clean, single-file “runner” structure, NERV empowers experienced users to directly customize the `RunTrials()` loop and individual state handlers, leveraging the full power of Unity’s `C#` scripting environment while still benefiting from NERV’s automated infrastructure for data management, hardware integration, and experiment execution. This balance between transparency and automation enables researchers to create highly customized experiments that meet their specific needs while preserving the rapid prototyping advantages of the no-code workflow.

### 2.4. Data Management & Archival

NERV builds on its transparent state-machine runner to capture the complete experimental context—code, configuration, screen images, and hardware pulses—into a self-contained, timestamped archive. At session launch a manifest header records system metadata and file checksums; during runtime the SessionLogManager streams frame-locked events into dual-clock CSVs, and upon completion it synthesizes a concise Task Summary to facilitate rapid quality control. The resulting folder hierarchy (Figure 6) provides both fine-grained records for detailed analysis and concise summary files for rapid quality control, ensuring that every experiment is reproducible and easy to audit.

**Figure 6:**
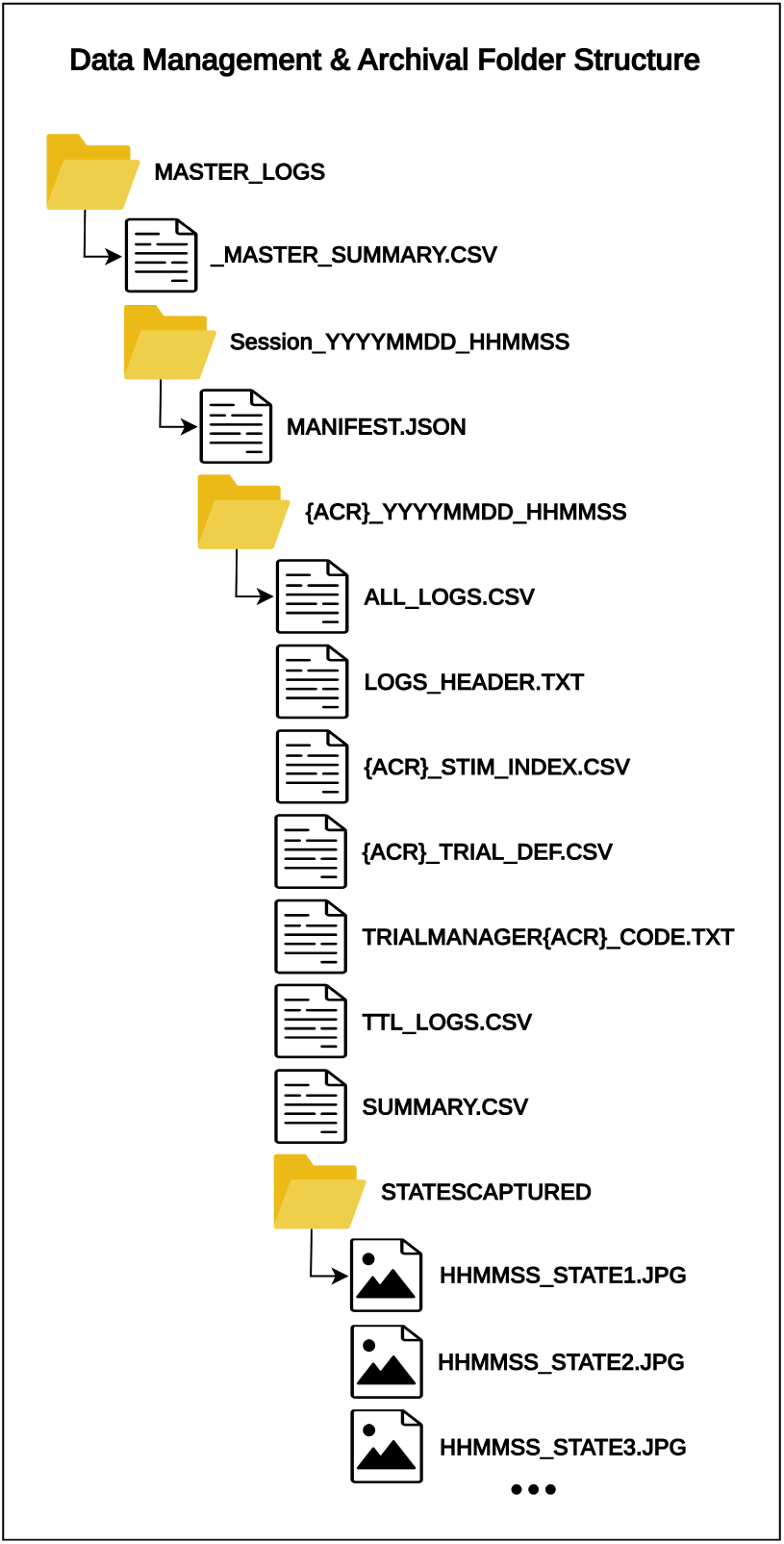
Data management folder hierarchy automatically generated by NERV. Each session is self-contained, with log files, configuration archives, and visual state captures to ensure reproducibility.

#### 2.4.1. Unified Session Manifest (MANIFEST.json)

The Unified Session Manifest (`MANIFEST.json`) is a cornerstone of NERV’s provenance strategy, encapsulating within a single, machine-readable file every datum necessary to reconstruct, audit, and re-execute an experimental session. Generated at launch, the manifest records dual timestamps (Unix epoch and ISO-8601 UTC) to support both automated parsing and human inspection; embeds a full system fingerprint including operating system build, GPU model, Unity version, display refresh rate, and the run-time `Application.targetFrameRate`; and binds the session to an immutable build identity via the Git commit hash injected during Unity build compilation. Crucially, for each task source code and configuration CSVs, a SHA-256 checksum is computed and stored. By coupling these cryptographic hashes with dual-clock event logging (Section 2.4.2), the manifest delivers bit-level reproducibility: every behavioral event can be traced unambiguously to the exact software state and stimulus set that generated it. Because the file is lightweight JSON, it functions simultaneously as a human-readable header for long-term archiving and as a drop-in metadata source for BIDS/NWB exporters, ensuring seamless interoperability with external toolchains. Collectively, this design bolsters trust, traceability, and longevity which are widely recognized as prerequisites for rigorous open science (Wilkinson et al., 2016; Nosek et al., 2015), by transforming each NERV session from a collection of log files into a a verifiable, tamper-resistant scientific record with full provenance. All manifest values can be seen in Table 1.

**Table 1:**
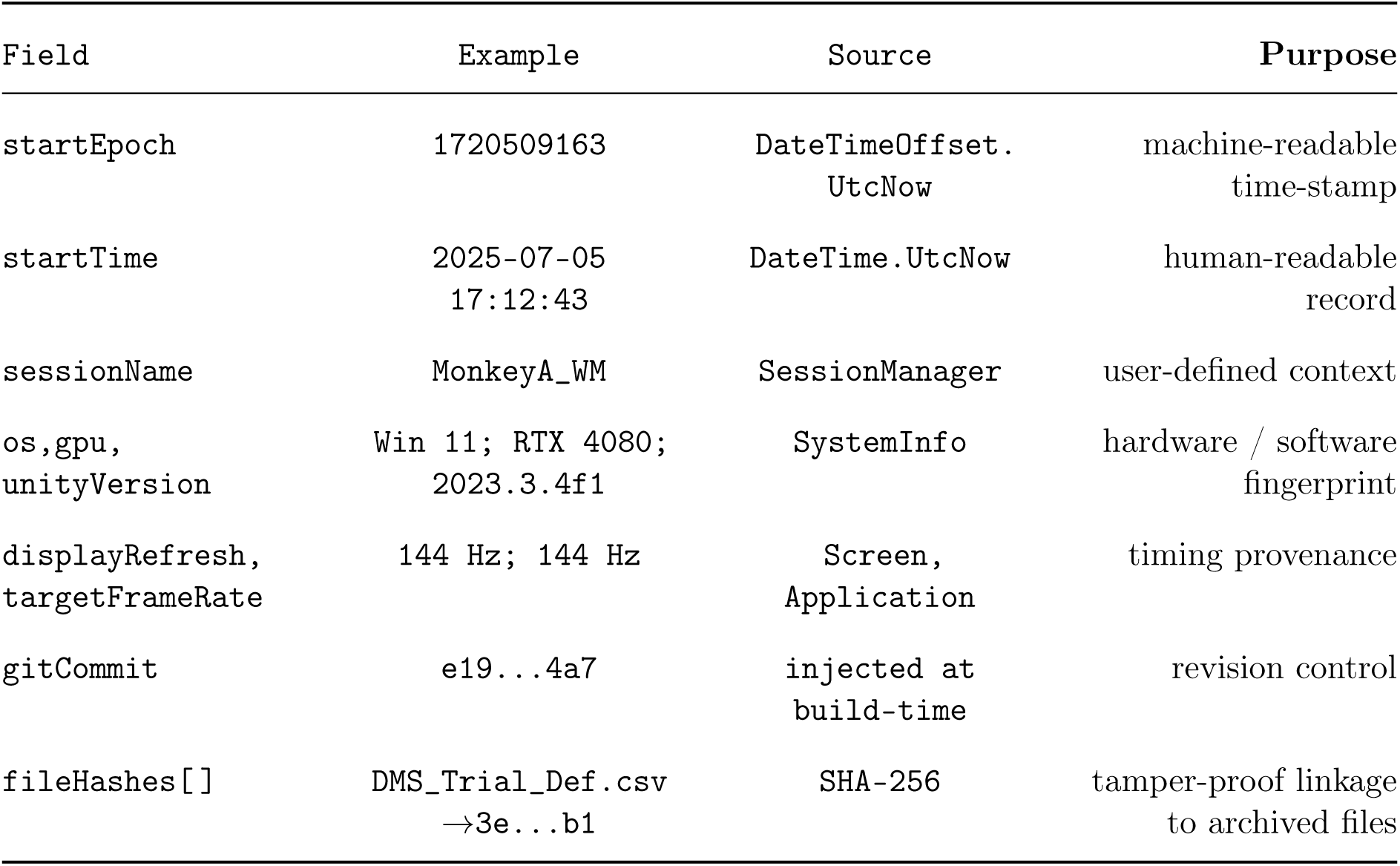
Session-level metadata recorded in MANIFEST.json.

#### 2.4.2. Runtime Logging and Data Archival

`SessionLogManager` is the backbone of runtime logging. Every behavioral and hardware event is time-stamped on two clocks, `Time.realtimeSinceStartup` (Unity) and `sw.Elapsed.TotalSeconds` (high-resolution CPU stopwatch), to enable millisecond-precision alignment with external recordings. All logs are stored under the project or build root directory in the `MASTER_LOGS` folder. The resulting archival folder hierarchy is summarized below:

- **_MASTER_SUMMARY.csv:** Aggregates task-level summaries across sessions into one file, providing a high-level overview of performance (Session Name, Task, Trials, Accuracy, Mean RT, Timestamp). Each task run appends a new row automatically (see Table 2).
- **Session Folder:** A timestamped folder (e.g., `Session_YYYYMMDD_HHMMSS`) containing all logs and data for a specific session.
- **MANIFEST.json:** A JSON file containing metadata about the session, including start time, system information, and checksums of all relevant files. This file serves as a comprehensive record of the session’s context and configuration (see Section 2.4.1).
- **Task Subfolder:** A subfolder created inside the session folder each time a specific task is started (e.g., `{ACRONYM}_YYYYMMDD_HHMMSS`).
- **LOGS_HEADER.txt:** Contains public parameters and settings from the TrialManager script for that run, providing a record of runtime configurations.
- **ALL_LOGS.csv:** Comprehensive event log with millisecond-resolution timestamps; each significant task event is written as a new row for post-hoc analysis. Entries are generated via the `LogEvent(“EventName”)` function within each `TrialManager` script. Columns include `Frame, UnityTime, StopwatchTime, TrialID, Event`, and Details.
- **TTL_LOGS.csv:** A specialized log that records the precise timing and code of all TTL pulse events sent to external hardware, enabling alignment with physiological recordings. Entries are generated via `LogEvent()` when the corresponding event is defined in the `TTLEventCodes` dictionary, initialized at the start of each `TrialManager` script. TTL event codes are assigned during the creation of the associated `ExperimentDefinition` asset. Columns include `Frame, UnityTime, StopwatchTime, TrialID, Event`, and Code.
- **StatesCaptured Subfolder:** Stores screenshots of the first user-defined occurrences of each unique trial state (e.g., the first two instances per state), enabling visual verification of the subject’s experience. Images are saved as `StateName_HHMMSS.png` at a user-defined resolution. Capture behavior and resolution can be configured in the SessionLogManager inspector.
- **SUMMARY.csv:** Task-level summary written at exit, recording aggregate metrics (Session, Task, Trials, Accuracy, Mean RT, Timestamp) along with per-trial results (TrialIndex, Correct, ReactionTimeMs).
- **TrialManager{Acronym}_CODE.txt:** A copy of the exact C# code for the TrialManager script used in that specific task run, ensuring full reproducibility and analysis consistency.
- **Experimenter_Comments.txt:** A text file containing all comments made in one of the dual display modes (e.g., `DualHuman` or `DualMonkey`) during the session, allowing experimenters to annotate the session with timestamped observations or notes (outlined in Section 2.5.2).
- **{Acronym}_Trial_Def.csv**: Archival copy of the exact trial-definition CSV for this run containing the full trial structure and condition definitions.
- **{Acronym}_Stim_Index.csv**: Archival copy of the indexed reference to all stimuli presented for this run.

**Table 2:**
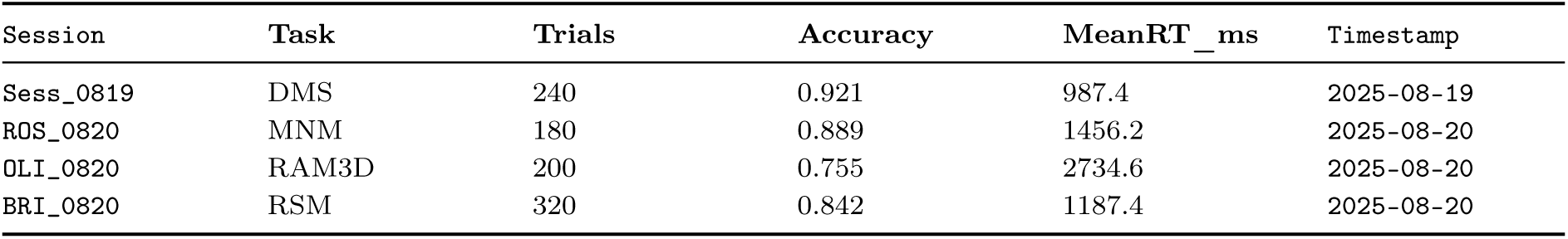
Example master summary rows (CSV structure preview)

This comprehensive archival system ensures full data provenance, allowing researchers or reviewers to reconstruct exactly what transpired in an experiment, from code execution to parameters, stimuli, and behavioral/system events.

### 2.5. Multi-Display Support and Experimenter Interface

To enhance experimenter oversight and minimize distractions for the subject during live experiments, NERV incorporates robust multi-display support, allowing for a dedicated subject display (Display 1) and a comprehensive experimenter interface on a second display (Display 2). This separation is crucial for maintaining experimental integrity while providing researchers with real-time monitoring and control capabilities.

NERV supports configurable display modes that can be dynamically loaded and saved at runtime, e.g. dedicating one display to a human experimenter controlling a monkey or human experiment:

- SingleHuman: A single display mode where all UI for both subject and experimenter interfaces are shown on Display 1. This is suitable for simple setups or when only one display is available.
- DualHuman: A dual-display mode where Display 1 is dedicated to the subject, showing only the task interface with blackout screens until the task screen is loaded, while Display 2 provides the experimenter with a comprehensive control panel, including real-time data monitoring, session management, and configuration options.
- DualMonkey: Similar to DualHuman, Display 1 is dedicated to the non-human primate subject, featuring blackout screens whenever the trial is not live, and minimal UI to prevent distraction, while Display 2 provides the experimenter with additional controls and monitoring tools.

These display modes are managed by the DisplayManager.cs singleton, which handles the activation of the appropriate displays and the visibility of UI elements based on the selected mode via the advanced settings menu within NERV (Figure 8B).

#### 2.5.1. Single Display Mode Overview (Display 1 only)

In Single Display Mode, all interfaces required for experiment setup and execution for both the subject and the experimenter are presented on Display 1. This includes setting the session name using the SessionNameInput field and selecting the desired task modules (experimental scenes) via toggles. Furthermore, communication port assignments for hardware integration (e.g., `COM3` for TTL via serial port) and the activation of a `testMode` can be configured through the `SettingsUIController` interface, accessible from the `TaskSelectorUI` (Figure 8A).

Once the experiment is launched, control of the interface transitions to the participant, eliminating the need for a secondary display. During actual task execution in `SingleHuman` mode (Figure 8C), NERV does not employ a blackout screen before the trial begins, in contrast to `DualHuman` and `DualMonkey` modes. While the `DualMonkey` mode hides most in-game UI to minimize distraction, in SingleHuman mode, essential UI elements such as the pause button, score display, feedback text, and other pause screen UI elements persist on Display 1 throughout the task.

It is important to note that the advanced experimenter monitoring and control features—such as the live subject preview with gaze visualization, the real-time debug log, live session statistics, on-screen variable editing, and the timestamped comment log—are specifically designed for and rendered on Display 2 in `DualHuman` and `DualMonkey` modes. Therefore, these dedicated experimenter tools are not available or visible during the subject’s task performance in `SingleHuman` mode to maintain a clean subject experience on the sole display.

#### 2.5.2. Dual Display Modes Overview (Display 1 and Display 2)

The dual display modes, `DualHuman` and `DualMonkey`, are designed to provide a comprehensive interface for both the subject and the experimenter, enhancing the experimental workflow and minimizing distractions for the subject. To achieve this, NERV utilizes a series of tailored blackout screens during experiment setup and transition states along with a dedicated experimenter interface on Display 2 to maximize focus and control.

**Display 1 (Subject View):** NERV’s multi-display modes provide tailored subject-facing behavior for each experimental context:

- DualHuman: Display 1 shows a blackout screen during setup and transitions between tasks, preventing the subject from viewing experimenter actions or sensitive information. Once a trial begins, the essential task interface is revealed, including the pause button, space for instructions, sound and settings buttons, and the score display. If enabled, the pause button, score, and feedback text persist throughout the entire task. All task input is restricted to Display 1, while Display 2 provides the experimenter with monitoring and configuration tools.
- DualMonkey: Display 1 remains blocked out during all non-task periods, including setup, inter-block intervals, and every pause. During active trials, only the minimal task view is shown, no UI elements, no score, and no words, to minimize distraction for non-human primate subjects. The screen returns to black immediately upon any pause or transition. Display 2 provides the same experimenter monitoring and configuration interface as in DualHuman mode.

**Display 2 (Experimenter Interface):** Display 2 provides the experimenter with a rich set of controls and real-time monitoring tools as seen in Figure 7. This includes:

1. Live preview of the subject’s screen, including gaze cursor overlay if eye-tracking is enabled.
2. Scrollable real-time event and debug logs for monitoring experiment flow, troubleshooting, and verification of hardware connections.
3. Session statistics and information such as accuracy, average reaction times (ms), labels for current/total trials and blocks, and elapsed time updated live.
4. On-screen controls for pausing, resuming, exiting, or editing task specific runtime variables.
5. Timestamped comment entry for annotating the session with experimenter notes saved to `SessionLogManager` task subfolder.

**Figure 7:**
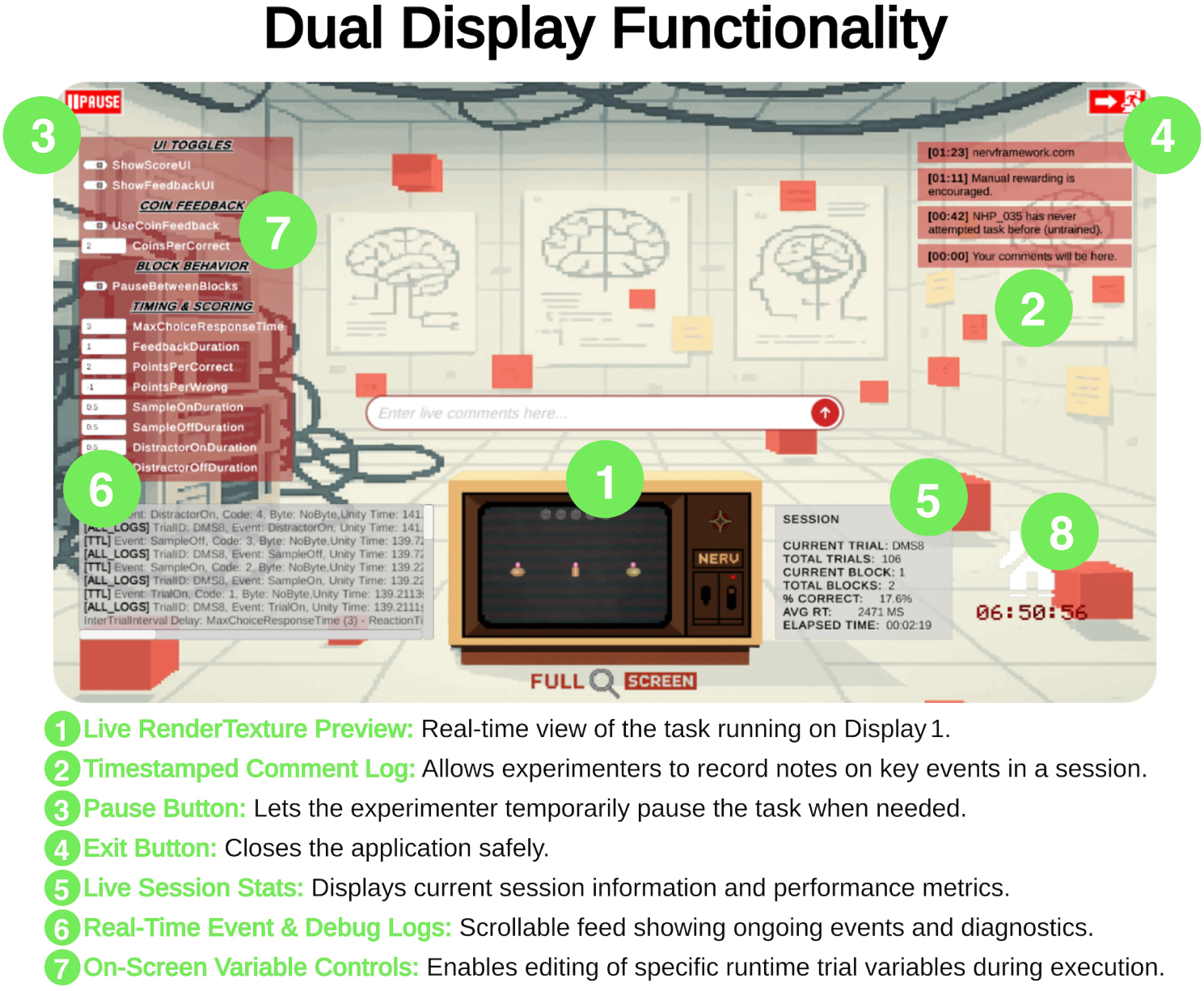
Experimenter-facing interface provided on Display 2 in NERV’s dual-display mode. The panel includes real-time task preview, logging, variable controls, and session monitoring tools, allowing experimenters to manage tasks and annotate sessions without disrupting the subject-facing display.

The experimenter interface can be enabled through the `Advanced Settings` panel of the the `TaskSelector` scene, which allows assignment of subject and experimenter displays. In this panel (Figure 8B), the user can configure one of the previously mentioned modes by selecting intuitive icons for each display. Display 2 is automatically configured to host the experimenter interface when applicable, delegating setup controls accordingly. These configuration settings are saved to `Application.persistentDataPath`, ensuring persistent customization across sessions. Once a task is loaded, the experimenter can monitor subject performance, manage task flow, and make real-time adjustments as needed.

**Figure 8:**
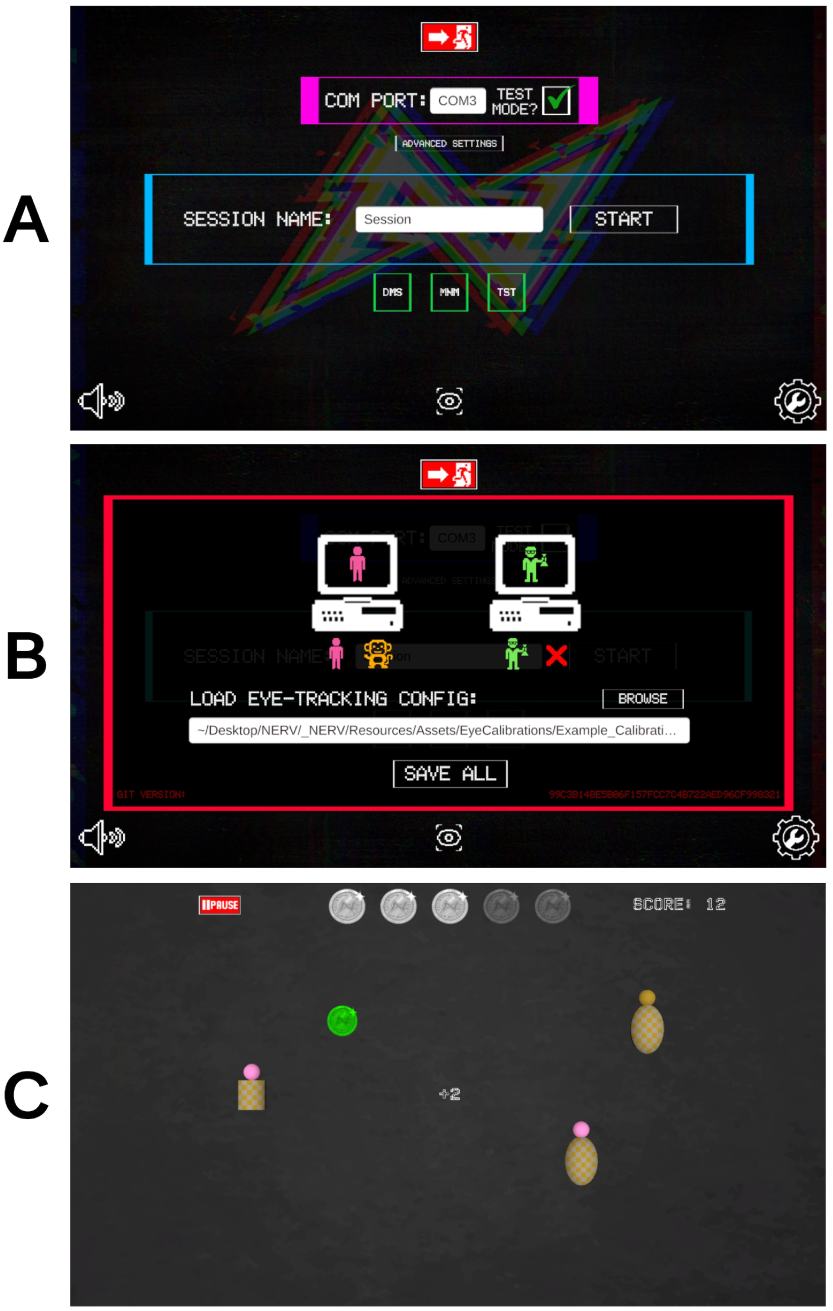
Graphical user interface for task selection, configuration, and execution. (A) **Task Selector UI.** Users specify session parameters (e.g., session name, COM port, test mode) and toggle the desired task from a menu (e.g., DMS, MNM, TST). (B) **Advanced Settings Panel.** Configuration screen for extended display modes and peripheral integration. Here, users load eye-tracking calibration files, enable dual-display functionality, and save global settings. (C) **In-game view (SingleHuman mode).** Example trial with real-time behavioral feedback, pause functionality, and score tracking displayed at the top of the screen, demonstrating how experimental stimuli and outcomes appear to participants.

### 2.6. Hardware Integration

NERV provides direct integration points with laboratory recording systems to ensure millisecond-level synchronization between behavioral events and physiological data. Current implementations support invasive neural recording setups (e.g., Neuropixels, SEEG) and eye-tracking via National Instruments NI-DAQ, while the same infrastructure can be adapted to other modalities as needed.

#### 2.6.1. TTL Output Hardware

Precise event-locked hardware synchronization is achieved in NERV via a USB-to-TTL interface built on the open-hardware design of Appelhoff and Stenner (2021). The interface exposes eight digital output lines driven by a micro-controller (Arduino Leonardo) that has been validated for millisecond latency and <1 ms jitter. Detailed build instructions and firmware are available online (https://stefanappelhoff.com/usb-to-ttl/index. html).

##### Unity side

At runtime, the singleton `SessionLogManager` opens a virtual COM port (default 115 200 bps) via `System.IO.Ports`. Whenever a state marked `IsTTL` begins, the manager writes a single byte to the port. Channel codes follow the bitmask convention 2*^n^*^−1^ for lines 1–7; TTL code 8 instead transmits `0xFF`, driving all eight lines high simultaneously. Each byte is time-stamped in `TTL_LOGS.csv`, ensuring hardware and behavioral logs share a common clock.

##### Micro-controller side

The Arduino firmware (unmodified from Appelhoff and Stenner (2021)) listens for the incoming byte, decodes the bitmask, and emits a 5 V TTL pulse (1 ms width) on the corresponding output pins. Because the decoding loop runs in interrupt-free polling at 8 MHz, turnaround time remains <200 µs on typical laboratory PCs.

##### Debugging mode

For bench testing or laptop demonstrations, a `Test Mode` toggle (available in the `SessionLogManager` Inspector and the on-screen settings panel of `TaskSelector.unity`) disables the serial handshake while still logging the intended pulses. This allows developers to verify task timing without connecting physical hardware.

The combination of Unity-level byte coding, a validated open-source TTL interface, and mirrored logging guarantees that every stimulus event delivered by NERV can be aligned with neural recordings to within millisecond precision.

For reproducibility, the full Arduino firmware that decodes the incoming byte and generates channel-specific TTL pulses is provided in the supplemental materials.

#### 2.6.2. Eye-Tracking Integration

NERV is designed to accommodate a range of eye-tracking hardware. In the current implementation, it supports analog input from devices such as ISCAN systems, interfaced through National Instruments (NI) data acquisition hardware. A custom `C#` wrapper interacts directly with NI’s `DAQ API` (`nicaiu.dll`) to stream real-time analog eye-position signals. The system includes a built-in calibration routine that converts raw voltages to gaze coordinates, facilitating accurate gaze visualization and gaze-contingent stimulus control. All DAQ interactions are centralized in `GazeCursorController.cs`, which reads analog inputs and utilizes `CalibrationLoader` to translate these voltages into on-screen gaze positions. The `CalibrationLoader` is instantiated automatically as part of the Dependencies prefab in every scene, so users do not need to manually add or configure it. By centralizing hardware communication and event logging, NERV ensures synchronized alignment between gaze data, behavioral events, and neural activity for streamlined analysis.

To implement a different eye-tracking system, a general integration workflow is as follows:

1. Install all required drivers and SDKs for the eye tracker.
2. Run the eye tracker’s calibration routine to map gaze data to screen coordinates.
3. Drag the `DwellClick.cs` ExtraFunctions script onto the `TaskManager{Acronym}` GameObject in your eye-tracking capable scene.
4. Edit the `DwellClick.cs` script to include the necessary API calls for your eye tracker, such as initializing the tracker, starting data collection, and retrieving gaze coordinates.
5. Map the `screenPos Vector2` coordinates to your eye tracker API’s output format, ensuring that gaze coordinates align with Unity screen space.
6. If visualization of the `GazeCursor` is desired, enable the corresponding option in the inspector of the `TrialManager{Acronym}` GameObject.
7. Test the integration by running a simple task and verifying that gaze data is captured correctly and stimuli can be selected in the “Choice” state of the task.

### 2.7. Example Experiments

NERV includes five example tasks that highlight its flexibility and provide a foundation for new users (Figure 9). These examples, available in the GitHub repository (https://github.com/kylecoutray/NERV) were implemented in Unity with development times ranging from minutes (e.g., simple DMS) to several hours (e.g., RAM3D). Collectively, they span classic 2D working memory paradigms, immersive 3D navigation tasks, and rule-based sequence manipulations, showcasing how the framework supports both rapid prototyping and more elaborate experiment designs.

- **Distractor Match-to-Sample (DMS):** Delayed matching with optional distractors during the delay interval.
- **Match/Non-Match (MNM):** Two-stimulus comparison assessing encoding and recognition.
- **Rule-Adaptive Match 3D (RAM3D):** 3D navigation requiring application of a context-dependent rule (match vs. non-match) during spatial choice.
- **Rule-Based Sequence Manipulation (RSM):** Sequence transformation requiring application of rule-based operations to maintained stimuli.
- **Sequence Distractor Match-to-Sample (SDMS):** Sequence comparison with distractor events introduced during the delay interval.

**Figure 9:**
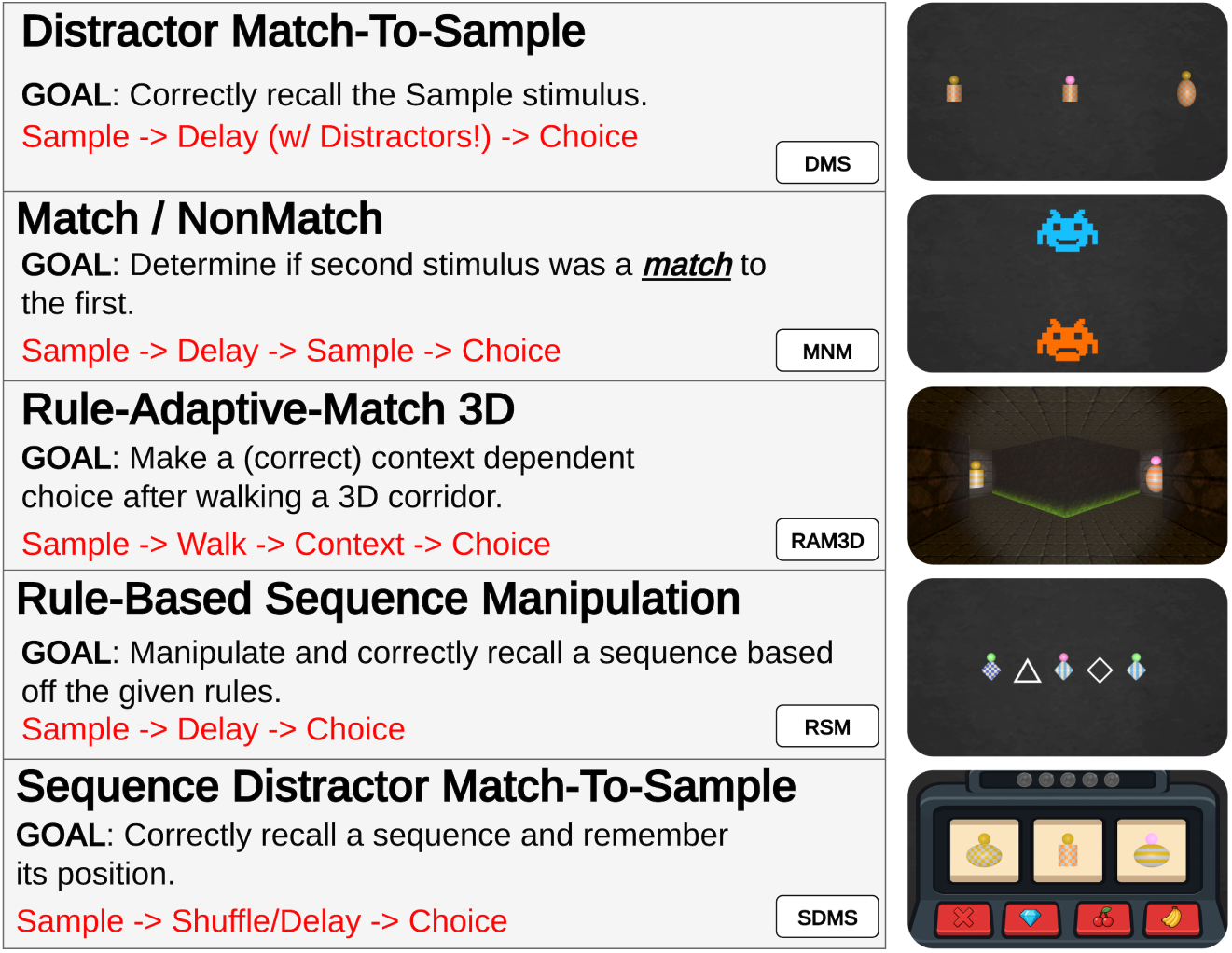
Overview of the five example experiments included in the NERV framework.

A live, web-based demonstration is available at https://nervframework.com and the open-source repository provides full implementations, including code structure and stimulus configurations.

## 3. Results

We evaluated NERV’s performance in terms of temporal precision, quantified through empirical measurements of latency and jitter across software, hardware, and display events. These results demonstrate that NERV preserves millisecond-level accuracy in event timing, ensuring reliable synchronization across behavioral and neural data streams.

### 3.1. Timing and Performance Validation

NERV is designed to achieve millisecond precision in event timing and maintain consistent frame rates, comparable to established toolkits such as Psychtoolbox (Brainard, 1997) and commercial platforms like Presentation (Neurobehavioral Systems, Inc., 2025). Leveraging Unity’s compiled architecture, latency and jitter for hardware synchronization events (e.g., TTL pulses, eye-tracker signals) remain exceptionally low.

#### Internal (Code-Level) Precision

Within Unity’s scripting pipeline, NERV executes critical events (stimulus instantiation, photodiode activation, CSV writes, TTL sends) within a single `Update/LateUpdate` cycle. This design ensures that NERV’s code-level events are deterministically aligned to one Unity frame, with `Time.frameCount` providing a stable temporal anchor for all software triggers.

To enforce this consistency, NERV enforces fixed Unity project settings in `SessionManager.cs`: `QualitySettings.vSyncCount` is set to 1 (vertical synchronization enabled), and `Application.targetFrameRate` is matched to `Screen.currentResolution.refreshRateRatio` (i.e., the display’s native refresh rate). This combination synchronizes buffer swaps with the display’s refresh cycle, preventing drift from asynchronous updates and ensuring consistent frame pacing (Garaizar et al., 2014).

#### External Hardware and Display Latency

Although Unity’s internal scheduling aligns all code-level events to single-frame precision, two external processes could, in principle, add delays: (1) GPU scanout to the display, and (2) serial driver latency in TTL transmission. However, because NERV enforces vertical synchronization and matches the frame rate to the display’s refresh rate, these delays are consistent and bounded (Elze, 2010). In practice, our empirical measurements (see below) demonstrate sub-frame jitter for screen updates and sub-millisecond jitter for TTL pulses, confirming that external latencies do not compromise millisecond-level synchronization.

#### Measurement and Time Alignment Methodology

To validate these guarantees, we simultaneously recorded visual and electrical outputs against NERV’s internal logs. Specifically:

1. **Code→Screen Latency:** Measured by capturing photodiode traces (via PhotodiodeMarker.cs) and comparing onset times with Unity log timestamps.
2. **Code→Pulse Latency:** Measured by recording TTL line outputs from the Arduino interface and comparing them to Unity log timestamps of pulse initiation.
3. **Offset Correction:** Mean latencies and jitter were computed for both paths, and trial logs were corrected by applying these offsets to align behavioral, visual, and neural events on a common clock.

All measurements were conducted on a Dell Latitude 7455 equipped with a Qualcomm Snapdragon X Elite SoC (X1E-80-100), integrated Adreno X1-85 GPU, 16 GB memory, and Windows 11. Stimuli were presented on the built-in 14” touchscreen display operating at 60 Hz.

To capture pixel onset and TTL events simultaneously, we affixed a photodiode to the display, connected the Arduino TTL output via COM port, and routed both signals into an Open Ephys acquisition board (v2.4, rev. 5). The setup is illustrated in Figure 10A.

**Figure 10:**
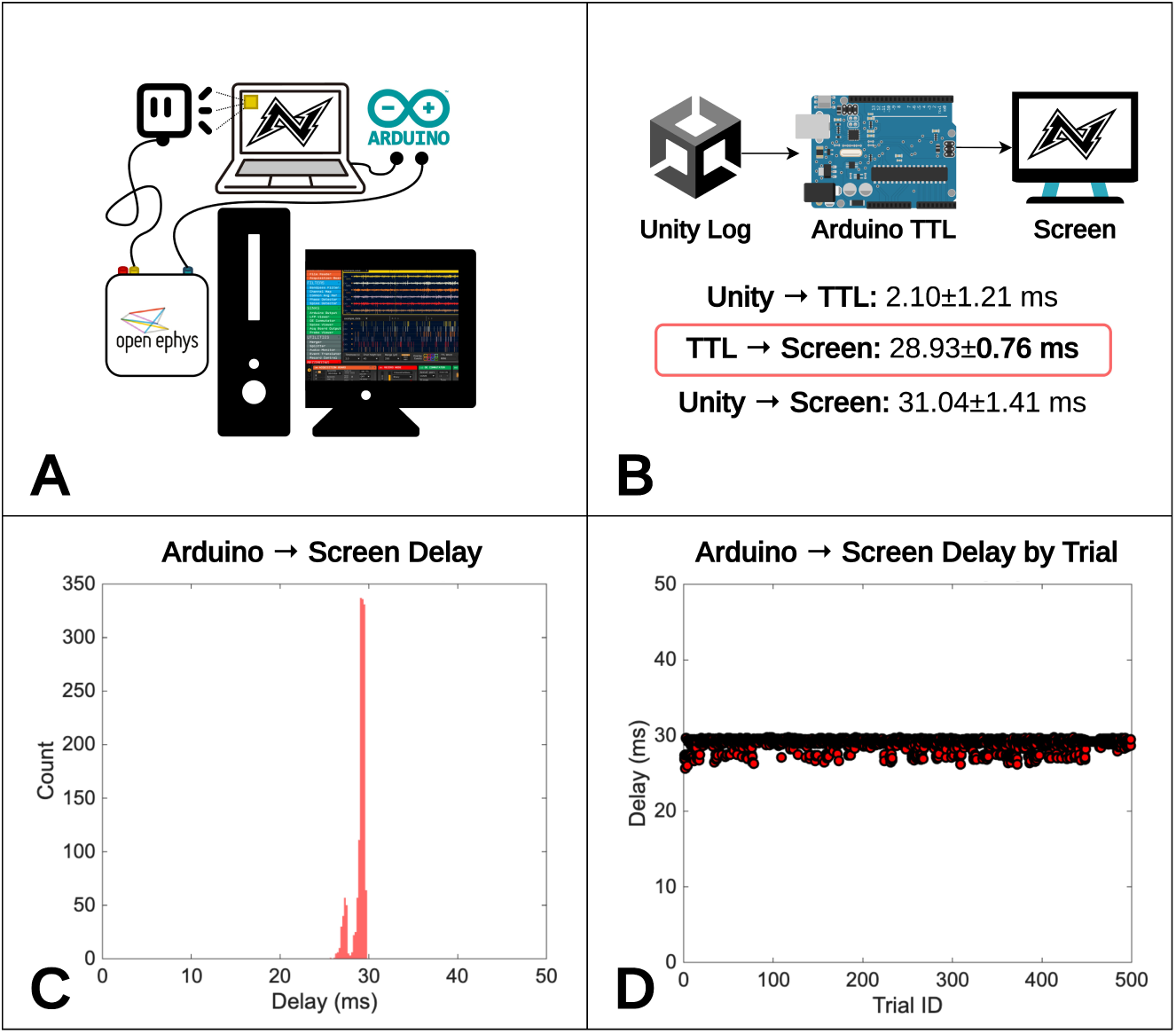
(A) Experimental setup integrating Unity for task control, Arduino for TTL pulse generation, and Open Ephys for pulse/photodiode signal acquisition. (B) Schematic of event timing: Unity log entries preceded TTL pulse generation by 2.10 ± 1.21 ms, while screen updates lagged the TTL pulse by 28.93 ± 0.76 ms. (C) Distribution of Arduino-to-screen delays across trials, showing a tight clustering around 29 ms. (D) Trial-by-trial delays demonstrating stable temporal alignment across 500 events.

#### Timing Validation Data

A 500-trial stress test confirmed NERV’s robust timing performance:

- **Software Delay (Unity → TTL): 2.10 ± 1.21 ms.** Derived from synchronizing Unity log timestamps with the physical TTL output, this primarily reflects jitter in the software-hardware interface and demonstrates precise control (Figure 10B).
- **Screen Delay (TTL → Photodiode): 28.93 ± 0.76 ms.** The interval between the TTL trigger and the first detectable pixel illumination, representing the key metric for aligning event logs with neural recordings (Figure 10C-D).
- **Total Delay (Unity → Screen): 31.04 ± 1.41 ms.** The combined end-to-end delay from Unity event log to visual stimulus appearance (Figure 10D).

These results demonstrate sub-millisecond jitter and highly consistent end-to-end latencies, confirming that NERV provides the temporal precision required for accurate alignment of neural recordings with behavioral events.

All raw data, detailed latency distributions (including TTL and eye-tracker events), and analysis code are provided in the supplemental materials to fully substantiate NERV’s timing performance.

## 4. Discussion

This study demonstrated that NERV enables experimenters to rapidly generate runnable tasks while preserving millisecond-level precision across software, hardware, and display events. These results position NERV as a unique contribution among existing experimental frameworks, uniting rapid development workflows with rigorous timing guarantees and comprehensive data provenance.

### 4.1. Comparison with Existing Tools

Relative to widely used packages, NERV addresses long-standing gaps by combining development efficiency, extensibility, and reproducibility in a single framework. Our laboratory has previously developed custom solutions for visual behavioral experimentation (Meyer and Constantinidis, 2005), which provided precise stimulus control but were tailored to specific paradigms and lacked the modularity needed to support diverse experimental designs and modern hardware integration. This experience directly motivated the development of NERV as a more generalizable, extensible framework.

Beyond our own tools, MATLAB/Psychtoolbox (Brainard, 1997) has long served as the reference standard for precise stimulus timing, but it requires substantial programming expertise and lacks integrated provenance tools. GUI builders such as PsychoPy (Peirce et al., 2019) and OpenSesame (Mathôt et al., 2012) lower the entry barrier for non-programmers but are constrained when extended to 3D environments or hardware-intensive tasks. Commercial options like Presentation (Neurobehavioral Systems, Inc., 2025) and E-Prime (Schneider et al., 2002) offer polished interfaces but remain closed-source and difficult to extend.

In contrast, NERV delivers millisecond-accurate timing comparable to Psychtoolbox while uniquely integrating a *no-code* pathway for rapid prototyping, a transparent scripting pathway for custom logic, and a Unity-based runtime capable of both 2D and immersive 3D experiments. Our evaluation confirms that this hybrid design combines precise temporal control with a workflow that enables rapid prototyping, positioning NERV as a next-generation platform for experimental neuroscience.

A key distinction is NERV’s transparent and extensible architecture. Whereas many GUI builders operate as *black boxes*, NERV generates a human-readable, well-commented `TrialManager` script. Researchers can directly modify trial logic, implement custom scoring algorithms, or create dynamic stimulus behaviors without bypassing the framework. Extensibility is further supported by the modular `ExtraFunctions` system, which uses event broadcasting to enable drag-and-drop components that can be added non-invasively to existing experiments. This design contrasts with the fixed element sets of traditional GUI tools, allowing new features to be incorporated flexibly and transparently.

Equally important is NERV’s integrated data management. At runtime, `SessionLogManager` automatically creates a timestamped folder hierarchy, archives the exact `TrialManager` source code and configuration CSVs, and records behavioral and hardware events (`ALL_LOGS.csv, TTL_LOGS.csv`) at millisecond resolution using both Unity’s monotonic clock and a high-resolution CPU stopwatch. Screenshots of displayed states are also stored via the `StatesCaptured` subfolder for visual verification. This strategy ensures full provenance: every experiment can be reconstructed exactly as it was run.

Finally, NERV centralizes experimental design, hardware interfacing, and data capture into a unified workflow. This reduces the fragmentation common in toolchains that combine separate stimulus, logging, and hardware systems, thereby minimizing transcription errors and enhancing reproducibility. By merging the engaging design potential of modern game engines, the accessibility of graphical builders, and the scientific rigor of precise timing and provenance, NERV provides a robust new platform for the neuroscience methods community.

### 4.2. Limitations and Future Directions

While NERV offers significant advantages, several practical considerations remain. Compared to lightweight script-based tools, Unity requires a modest initial setup, and although coding is unnecessary for standard experiments, basic familiarity with the engine can accelerate adoption. Predefined trial state types (e.g., `IsStimulus, IsDelay, IsChoice, IsFeedback, IsClear`) enable rapid construction of straightforward paradigms in minutes, whereas complex tasks with custom logic still demand engagement with Unity’s scripting environment.

Hardware integration is another area for expansion. Current implementations (TTL signaling and ISCAN eye-tracking) meet our labratory’s immediate needs, but other experimental setups may require additional customization. Future work will focus on expanding compatibility (e.g., joystick controllers, alternative DAQs) and releasing comprehensive documentation and tutorials to reduce barriers for new users.

As an open-source project, NERV is designed to evolve. The modular `ExtraFunctions` architecture facilitates community-driven contributions, enabling new experimental features and hardware interfaces to be added without disrupting the existing framework. The official GitHub repository serves as the central hub for updates, collaborative efforts, and community support, ensuring that NERV remains adaptive and continues to advance the design of neuroscience experiments.

### 4.3. Conclusion

NERV provides a unified, open-source framework that integrates rapid prototyping, rigorous temporal precision, and comprehensive data provenance for neuroscience experimentation. By embedding experiment logic within the Unity engine, it combines the accessibility of graphical builders with the extensibility of full C# scripting, offering a *“low floor, high ceiling”* design adaptable to both novice and expert users.

Empirical validation confirmed that NERV achieves millisecond-level timing precision across software, hardware, and display events, meeting the stringent requirements for integration with neural recording systems. The framework’s automated archival of code, configuration files, logs, and visual records ensures reproducibility and long-term verifiability, transforming each experimental run into a complete scientific record. The example tasks demonstrate that a wide range of tasks—from canonical match-to-sample designs to immersive 3D rule-based navigation—can be prototyped and executed within hours, underscoring both the efficiency and flexibility of the system.

Looking forward, NERV’s modular design and open-source availability create a foundation for community-driven development. By facilitating the integration of new task logic, hardware interfaces, and analytical pipelines, NERV offers a scalable solution that unifies stimulus control, hardware synchronization, and reproducible data management. In doing so, it not only lowers barriers to innovation but also establishes a rigorous, extensible foundation for the next generation of neuroscience experimentation.

## Acknowledgements

We thank Zhengyang Wang for his generous help with lab infrastructure questions and Peiyu Chen for valuable comments on an earlier version of the manuscript. This work was supported by the VUSE Summer Research Program and NIH grant R01 EY017077.

## Code Availability

NERV is released as open-source under the MIT licence, and the full source code, documentation, and example tasks are available at https://github.com/kylecoutray/NERV

